# A physiologically-validated rat model of term birth asphyxia with seizure generation after, not during, brain hypoxia

**DOI:** 10.1101/2020.05.05.078220

**Authors:** Tommi Ala-Kurikka, Alexey Pospelov, Milla Summanen, Aleksander Alafuzoff, Samu Kurki, Juha Voipio, Kai Kaila

## Abstract

Birth asphyxia (BA) is often associated with seizures which emerge during the recovery and may exacerbate the ensuing hypoxic-ischemic encephalopathy. In rodent models of BA, exposure to hypoxia is used to evoke seizures, which commence already during the insult. Here, we introduce a term-equivalent model of BA, in which seizures are triggered after, not during, brain hypoxia. Postnatal day 11-12 rat pups were exposed either to steady asphyxia (15 min; 5 % O_2_ + 20 % CO_2_) or to intermittent asphyxia (30 min; three 5+5 min cycles of 9 % and 5 % O_2_ at constant 20 % CO_2_). Cortical activity and seizures were recorded in freely-behaving animals. Simultaneous electrode measurements of cortical local field potentials (LFP) and intracortical pH and *P*o_2_ were made under urethane-anesthesia. Both protocols decreased blood pH to <7.0 and base excess by 20 mmol/l, and evoked an increase in plasma copeptin (0.2 to 5 nM). Clonic and tonic convulsions were triggered after intermittent but not steady asphyxia, and they were tightly associated with electrographic seizures. During intermittent asphyxia LFP activity was suppressed as brain pH decreased from 7.3 to 6.7. Brain *P*o_2_ fell below detection level in 5 % ambient O_2_ but returned to the baseline level during steps to 9 % O_2_. Neuronal hyperexcitability and seizures were suppressed in all types of experiments when the post-asphyxia brain pH recovery was slowed down by 5 % CO_2_. Our data suggest that the recurring hypoxic episodes during intermittent asphyxia promote neuronal excitability, which becomes established as hyperexcitability and seizures only after the suppressing effect of the hypercapnic acidosis is relieved. The present rodent model of BA is to our knowledge the first one in which, consistent with clinical BA, robust behavioral and electrographic seizures are triggered after and not during the BA-mimicking insult.

## INTRODUCTION

Birth asphyxia is one of the leading causes of neonatal mortality resulting in around one million neonatal deaths annually (Lawn et al., 2005). In survivors, BA leads to hypoxic-ischemic encephalopathy (HIE) making them prone to a wide variety of developmental aberrations and lifelong malfunctions of the brain, ranging from minor and major psychiatric and neurological disorders to cerebral palsy (Ahearne et al., 2016; Dalman et al., 2001; Modabbernia et al., 2016; Pappas and Korzeniewski, 2016; Rosso et al., 2000). The period of recovery after asphyxia is often marked by seizures, some of which are considered to promote HIE-related trauma (Bennet et al., 2007; Glass et al., 2009; Kharoshankaya et al., 2016; McBride et al., 2000; Miller et al., 2002; Murray et al., 2009; Pressler and Mangum, 2013; Soul et al., 2019; van Rooij et al., 2010; Wirrell et al., 2001).

By definition, asphyxia is a combination of systemic hypoxia and hypercapnia, and these two components of asphyxia have distinct – often functionally opposite – actions on the brain. Hypoxia is known to promote neuronal excitability and seizures (Jensen et al., 1991; Kawasaki et al., 1990; Peng et al., 2013; Sampath et al., 2014; Zanelli et al., 2014 and references below), whereas an elevation of systemic CO_2_ produces a fall in brain pH and a consequent decrease in neuronal excitability (Lee et al., 1996; Pasternack et al., 1996; Ruusuvuori and Kaila, 2014; Schuchmann et al., 2006; Shi et al., 2017; Tolner et al., 2011). Moreover, hypercapnia leads to vasodilation of cerebral arteries and arterioles (Giussani, 2016; Vutskits, 2014) and, acting in synergy with neurohormonal factors such as vasopressin (Perez et al., 1989), mediates a brain-sparing increase in cerebral blood flow and oxygenation during the asphyxia-coupled reduction of O_2_ availability (Giussani, 2016).

Despite the differences in the neurobiological and physiological effects of hypoxia and asphyxia, practically all models on BA employing neonatal rodents are based on exposure of the animals to hypoxia or hypoxia-ischemia (Jensen et al., 1991; Rice et al., 1981; Sun et al., 2016; Vannucci and Vannucci, 2005). In such models, seizures are triggered *during* the exposure to hypoxia (Jensen et al., 1991; Sampath et al., 2014). This is in contrast with observations of human neonates in which seizures are triggered *after* a period of moderate or severe asphyxia (Lynch et al., 2012).

In order to analyze the fundamental differences between hypoxia- and (post-)asphyxia-induced seizures, we investigated the dependence of seizure generation on changes in brain pH/CO_2_ and oxygen levels. The present rat model was recently used to study hormonal and brain-protective responses in asphyxiated rodents (Pospelov et al., 2020; Summanen et al., 2018). Here we used postnatal-day (P) 11-12 rats which, in terms of cortical development, are at a stage that is equivalent to the human term neonate (Clancy et al., 2007; Romijn et al., 1991; Semple et al., 2013). Asphyxia was induced by ambient gas mixtures containing 5 % or 9 % O_2_ and 20 % CO_2_ (balanced with N_2_). Two protocols were used in the present study: *steady asphyxia* (5 % O_2_ plus 20 % CO_2_) which mimics an acute complication such as placental abruption or maintained cord compression; and *intermittent asphyxia* where the hypoxia is applied in repetitive steps at 9 % O_2_ and 5 % O_2_ (at constant 20 % CO_2_) in order to roughly mimic the effects of recurring contractions during prolonged parturition.

In sharp contrast to BA models based on exposure to pure hypoxia, seizures were not observed during the profound brain hypoxia which takes place during asphyxia with 5 % O_2_. However, intense behavioral convulsions, tightly paralleled by electrographic cortical seizures, were triggered during brain pH recovery after the intermittent asphyxia protocol. The seizures were strongly suppressed when the rate of brain pH recovery was slowed down by a low level (5 %) of ambient CO_2_. No seizures were seen after steady asphyxia. The striking difference in seizure propensity following the steady and intermittent asphyxia protocols suggests that periodic hypoxic episodes in the asphyxic brain enhance neuronal excitability (Jensen and Wang, 1996; Quintana et al., 2015; Zanelli et al., 2015), which becomes established as hyperexcitability and seizures once the suppressing effect of the asphyxia-related hypercapnic acidosis is relieved. Our data as a whole are consistent with the fact that human neonates are typically prone to seizures *after*, but not during parturition.

## MATERIALS AND METHODS

### Animals

Experiments were performed using postnatal day (P) 11-12 male Wistar Han rats (*n* = 166). The animals were maintained in an in-house animal facility under 12-hour light/dark cycle (lights on at 6 am) with a temperature and relative humidity of 21-23 °C and 45-55 %, respectively. Water and food (Altromin 1314 Forti, Lage, Germany) was available *ad libitum*. All experiments were carried out in accordance with the ARRIVE guidelines and the European Union Directive 2010/63/EU on the protection of animals used for scientific purposes. They had been approved by the National Animal Ethics Committee of Finland (license: ESAVI/6135/04.10.07/2015), and the Animal Ethics Committee of the University of Helsinki (license: KEK15-013). A maximum of two animals per litter were used, and littermates were never included in a given experimental group.

### Experimental asphyxia

The rat pups were placed in a 2000 ml chamber, with a constant temperature of 36 °C, 30-45 min prior to exposure to asphyxia. All gases were humidified and warmed, and delivered to the chamber at 2000 ml/min. Under these conditions the animals’ rectal temperature stabilizes to 36.5 – 37.0 °C. The gas mixtures were either provided by AGA (Espoo, Finland) or made on-site using an Environics S4000 gas mixing system (Tolland, Connecticut, USA).

Two distinct asphyxia protocols, steady asphyxia and intermittent asphyxia, were used to simulate BA (see Introduction). In steady asphyxia, the animals were exposed to 5 % O_2_ / 20 % CO_2_ (balanced with N_2_) for 15 min after which they were promptly re-exposed to room air and monitored for up to 90 min. The intermittent asphyxia consisted of three 5+5 min cycles of 9 % and 5 % O_2_ at constant 20 % CO_2_ with a total duration of 30 min (see Fig. 1). After the intermittent asphyxia, the first 30 min of recovery occurred either in room air (rapid restoration of normocapnia, RRN) or in 21 % O_2_ / 5 % CO_2_ gas (graded restoration of normocapnia, GRN) and was followed by 60 min in room air. Rats in the control group were kept in the experimental chamber, in room air, for an equivalent time.

**Fig 1.**
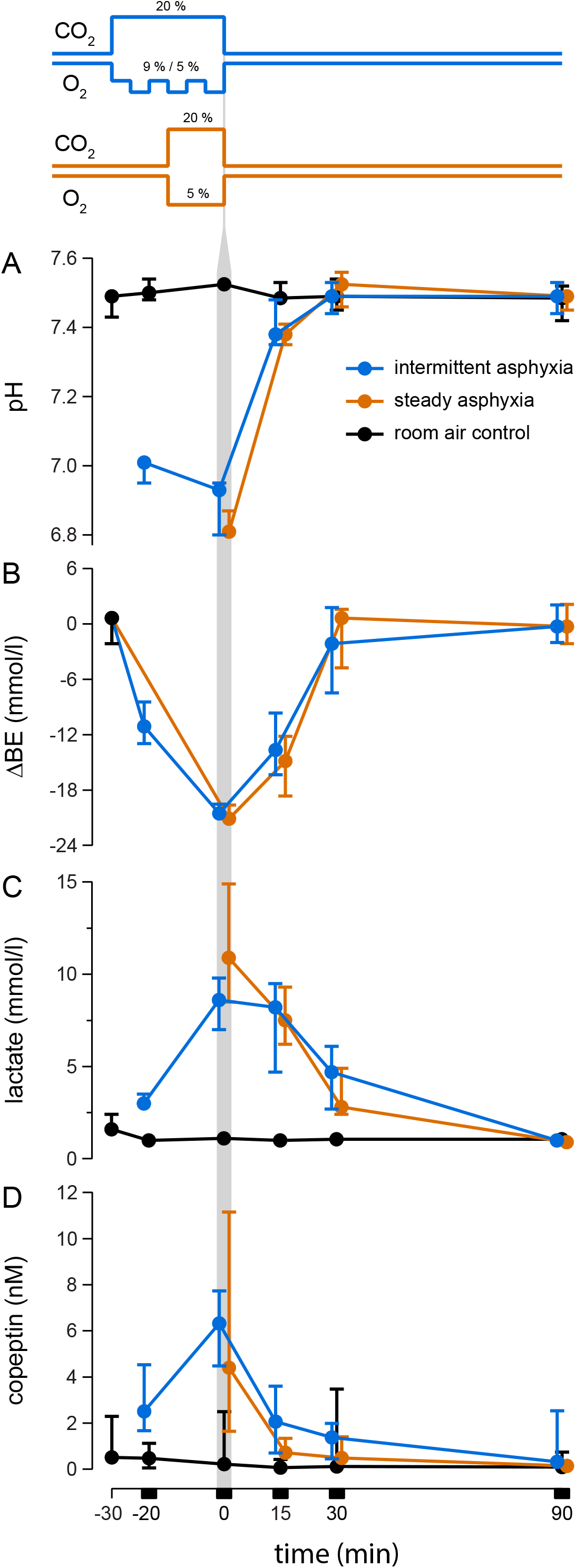
Changes in blood pH **(A)**, base excess **(B)** lactate **(C)**, and plasma copeptin **(D)** in P11-12 rats exposed to experimental asphyxia. The line graphs on the top illustrate the intermittent and steady asphyxia protocols. Both protocols decreased pH to < 7.0 and BE by 20 mmol/l (ΔBE), and evoked a marked increase in lactate and copeptin. The vertical grey bar marks the end of asphyxia, and, for clarity, the blue symbols have been shifted slightly to the left and the brown symbols slightly to the right. The values are median with 95 % CIs. All data are based on P11-P12 rats in this and subsequent figures.

### Blood sample collection

The animals were decapitated, and 80 μl of blood was collected straight from the trunk into lithium heparin coated plastic capillaries for blood-gas and lactate measurements. The rest of the trunk blood was collected into EDTA-coated tubes with protease inhibitors (Complete, Roche). The samples were centrifuged 10 min at 1300 g in 4 °C. Plasma samples were stored at −80 °C for copeptin analyses.

### Blood-gas and copeptin measurements

Blood pH, *P*co_2_ and lactate were measured using a GEM Premier 4000 Blood-gas analyzer (Instrumentation Laboratory, Lexington, Massachusetts, USA). Base excess (BE) was calculated from pH and *P*co_2_ as described in Summanen *et al.* (2018). The analyzer is designed to detect pH ≥ 6.8 and *P*co_2_ ≤ 20 kPa. 1/5 and 2/5 animals sampled after intermittent and steady asphyxia, respectively, had blood pH lower and a *P*co_2_ higher than these limits. For calculating BE, pH was assumed in these cases to be 6.8 and *P*co_2_ to be 20 kPa. Thus, the calculated BE was −11.2 mmol/l. Notably, the median values given in the Results were not affected by this approximation.

Copeptin concentrations were measured using an assay based on AlphaLISA (PerkinElmer) as previously described (Summanen et al., 2018). In brief, 5 μl of blank, sample or standard (0-160 ng/ml) solution was pipetted in triplicate on a 384-well AlphaPlate. 10 μl of polyclonal goat anti-copeptin (sc-7812, SantaCruz Biotechnology) conjugated to AlphaLISA acceptor beads (total concentration: 20 μg/ml) was added to each well. The plate was incubated for 1 h before adding 10 μl of 5 nM biotinylated, polyclonal sheep anti-copeptin (a gift from ThermoFisher) and, after another 1 h incubation, 25 μl of streptavidin-coated AlphaLISA donor beads (40 μg/ml) to each well. After a final 1 h incubation, the plate was read with an EnSpire Alpha plate reader (PerkinElmer). The standard curve was calculated with GraphPad Prism software (San Diego, California, USA) using 4-parameter logistic regression.

### Behavioral seizure scoring

Animals were recorded during the experiments using either a Sony HDR-CX410 or a Logitech C310 video camera for off-line analysis of behavioral seizures. Seizures were graded using a modified Racine scale (Racine, 1972) consisting of Racine stages (RS) III, IV and V, which are considered to reflect the propagation of seizures from forebrain structures (RSIII) to the brainstem (RSV) (Kellaway and Hrachovy, 1983; Pospelov et al., 2016). The stages were defined as follows: (RSIII) uni- or bilateral forelimb clonus, (RSIV) uni- or bilateral forelimb clonus paralleled by loss of righting reflex and (RSV) tonus-clonus with loss of righting reflex. To qualify for a clonic seizure, unequivocal and uninterrupted clonic jerks with at least 5 repetitions had to be observed. Tonus-clonus was defined as tonic extension of the forelimbs and/or hindlimbs following clonic seizures or tonic forelimb extension coupled with hindlimb clonus. The observed tonus typically resembled decerebrate rigidity and often co-occurred with opisthotonus. During seizure grading two reviewers, blind to the treatment, recorded latency and duration of each seizure stage. The reported values were agreed on by both reviewers: when disagreements arose, a consensus was reached by discussion.

During the first 10 min after asphyxia, a number of behaviors that were not commonly seen in control animals were observed. These included imbalance, startles, shaking, swimming, pedaling, excessive grooming, kicking, chewing, Straub tail and circling. Some of these behaviors were likely non-convulsive seizures belonging to RSI (oral automatisms) and II (myoclonus). All of them were short-lasting, albeit often repetitive, and their presentation varied, which made their rigorous categorization difficult. Moreover, in human neonates myoclonus and automatisms generally do not have an electrographic correlate, and treating them as seizures is advised against (Boylan et al., 2015; Mizrahi and Kellaway, 1987; Pressler, 2012). Thus, we decided not to include them into the present study which characterizes our BA seizure model.

### Video-electrocorticography in freely moving animals

P10-11 rat pups were exposed to 4 % isoflurane in room air, and after the induction of anesthesia, isoflurane was reduced to 1.5-2.5 % and the animals were given 5 mg/kg karprofen (Rimadyl Vet, Pfizer) subcutaneously. The scalp was scrubbed with 100 mg/ml solution of povidone-iodine (Betadine, Takeda) followed by 70 % ethanol before applying 10 % lidocaine solution. A piece of the scalp was removed and cranial windows were drilled over the parietal (A/P −3.8, M/L 2.0 mm from bregma) and the frontal cortex (A/P +1.5, M/L 1.5 mm) for recording electrodes and over the cerebellum (approx. 1 mm caudal from lambda) for a ground electrode. Care was taken to avoid damaging the dura during the drilling. In a few (5/27) recordings, a second window was drilled over the cerebellum for a reference electrode. 0.8 mm steel screws soldered to a strip connector were placed over the dura. The screws, the connector, the wires and the exposed skull were covered with dental cement. After 30 minutes of recovery on a heating pad, the animals were returned to the litter.

On the day following the surgery, the pups were placed in the asphyxia chamber. A piezo sensor (AB1070B-LW100-R, PUI Audio, Dayton, Ohio, USA) was taped over the flank for recording movement and breathing rhythm. The movement recordings showed a very low contamination of the ECoG recordings by movement artifacts (see below, and SVideo 4). The electrocorticogram (ECoG) was recorded using extracellular field potential amplifiers (npi EXT-02F/2 [bandwidth 3 – 1300 Hz] and EXT-16DX [0.1 – 500 Hz]; Tamm, Germany). ECoG and piezo signals were digitized at 2 kHz with a Cambridge Electronic Design Micro1401-3 converter (Cambridge, United Kingdom). Most (22/27) recordings were referenced to ground while few were referenced to the cerebellar reference electrode. After 45 min of baseline recording, the animal was exposed to intermittent asphyxia.

### Brain pH/PO_2_ and local field potential recordings

In experiments with intracranial recordings, the rat pups were anesthetized with 4 % isoflurane in room air and 1 mg/g urethane was administered intraperitoneally. The animals were transferred on a 35 °C heating plate and isoflurane was reduced to 1.5 – 2.5 % for the surgery. A piece of the scalp was removed and cranial windows were drilled over the parietal cortex (A/P −4.5, M/L 2.5 mm from bregma) for a glass capillary local field potential (LFP) electrode (150 mM NaCl) and over the cerebellum for a Ag/AgCl ground electrode. Additional windows were drilled bilaterally over the parietal cortex (A/P −4.0, M/L 2.0 mm) for pH and *P*o_2_ sensitive microelectrodes (Unisense OX-10 and pH-25, Aarhus, Denmark). To aid fixing the animal, a plastic washer was attached to the skull with dental cement.

After surgery, isoflurane administration was terminated. The animal was transferred to the recording setup, placed on a heating pad and fixed from the head to a stereotaxic bench. Small-rodent facemask (model OC-MFM, World Precision Instruments, Sarasota, Florida, USA) delivering room air was placed lightly over the snout and a piezo sensor (PMS20, Medifactory, Heerlen, The Netherlands) was taped over the ribcage to record movement and breathing rhythm. Small incisions were cut to the dura for the pH, *P*o_2_ and LFP electrodes and the electrodes were lowered 1 mm into the cerebral cortex. The ground wire was placed over the cerebellum. Rectal temperature was monitored using an RET-4/RET-5 probe and a BAT-12 thermometer (Physitemp, New Jersey, New York, USA) and the temperature of the heating pad was adjusted to maintain rectal temperature at 36 °C. If the animal reacted to a light tail pinch during the setup procedure, an additional 0.25 – 0.5 mg/g urethane was given intraperitoneally.

Details of the recording setup and calibration of pH and *P*o_2_ probes are described in (Pospelov et al., 2020). After the pH and *P*o_2_ recordings and the animal’s rectal temperature had stabilized, 30-60 min after the termination of the isoflurane anesthesia, the recording was started. Simultaneous recordings of LFP, pH and *P*o_2_ were done in a total of 29 animals: LFP and *P*o_2_ were recorded from 16 and 17 animals, respectively, in the RRN group and 12 animals in the GRN group, whereas pH was available from 14 animals in the RRN group and 7 animals in the GRN group. To make the recordings comparable to the freely-moving experiments, a dead volume was added in front of the facemask and the gas flow was adjusted to match the slower rate of gas exchange in the chamber using an oximeter (TR250Z 25% oxygen sensor from CO_2_Meter, Inc. Ormond Beach, Florida USA).

### Data analysis and statistics

Electrographic seizures were manually annotated on bandpass filtered (3-40 Hz) ECoG and LFP recordings. Seizures were defined as abnormal, repetitive spike discharges with an amplitude exceeding baseline mean+5*[standard deviation] and with a duration of ≥10 s. Consistent spike bursts (even if shorter than 10 s) occurring within less than 10 s before or after the seizure epoch were considered to belong to the same epoch. Annotated electrographic and behavioral seizures were time-synchronized in Spike2 software v9.08 (Cambridge Electronic Design) according to a test light stimulus in the video.

ECoG and LFP power in the 3-40 Hz band was calculated in 30 s windows, adjacent windows separated by 3 s, and normalized to average baseline power. The pH and *P*o_2_ recordings were analyzed using custom-made Matlab (MathWorks inc.) scripts as previously described (Pospelov et al., 2020).

The statistical analyses were made using Prism v8.0.1 software (Graphpad, San Diego, California, USA) and R v3.6.1. The reported values are median with either 95 % confidence interval (CI) of median or range. The type of variation is stated on each occasion in the Results. Binary and continuous variables were compared using Barnard’s test (Lydersen et al., 2009) and Mann-Whitney U-test, respectively. The difference between groups was considered significant when two-tailed p-value was ≤0.05.

## RESULTS

### Changes in blood pH, base excess and lactate and plasma copeptin evoked by experimental asphyxia

To ascertain the translational relevance of the experimental models used in this work, and to compare the hypoxic burden brought about by the steady and intermittent asphyxia protocols, we measured blood gas parameters which are used in the diagnosis of BA. As expected (see Pospelov et al., 2020; Summanen et al., 2018), both types of asphyxia induced a large decrease in blood pH and base excess (BE) (Fig. 1 A-B). Median pH was 6.81 [95 % confidence interval (CI): 6.80-6.87] and 6.93 [6.80-6.95], respectively, at the end of steady and intermittent asphyxia; and in the control group at the same time point, 7.53 [7.51-7.54]. The acidemia had a prominent metabolic component, as indicated by a fall in BE of 21.2 [18.9-22.3] mmol/l and 20.6 [19.3-21.2] mmol/l during steady and intermittent protocol, respectively. These BE changes were closely paralleled by an increase in blood lactate from a baseline of 1.1 [1.0-1.2] mmol/l under control conditions to 10.9 [8.5-14.9] mmol/l and 8.6 [7.0-9.8] mmol/l for steady and intermittent asphyxia, respectively (Fig. 1C). In addition, the levels of plasma copeptin, a relevant biomarker of asphyxia (Evers and Wellmann, 2016; Kelen et al., 2017; Schlapbach et al., 2011), were highly elevated by the end of both asphyxia protocols (4.41 [1.64-11.15] nM and 6.32 [4.48-7.74] nM for steady and intermittent, respectively vs. 0.22 [0.1-2.5] nM in the control group; Fig 1D).

While the duration of the intermittent asphyxia exposure was twice as long as that of steady asphyxia, the blood gas parameters showed quantitatively similar changes. As described in detail before (Pospelov et al., 2020), rat pups have a remarkable ability to physiologically compensate in response to the 9 % O_2_ challenge. The near-identical hypoxic burden caused by the two asphyxia protocols is therefore readily explained by the identical total time (15 min) of exposure to 5 % O_2_. This similarity is particularly interesting when comparing the efficacy of steady and intermittent asphyxia to induce post-asphyxia seizures, as done below.

### Behavioral seizures emerge following intermittent asphyxia, and they are suppressed by Graded Restoration of Normocapnia

We started our experiments on seizure generation in the two asphyxia paradigms using freely moving, non-instrumented rats. Seizure scoring was based on a modified Racine scale (see Methods and Racine, 1972). Under control conditions, the pups were for most of the time in apparent sleep. During the exposure to steady or to intermittent asphyxia, the pups initially displayed distress behavior with increased locomotion, but no seizures were observed. Breathing rate decreased gradually and led to apnea and death in 3/17 and 2/17 pups during exposure to steady and intermittent asphyxia, respectively. There was no further mortality.

Notably, convulsive seizures (RSIII-V) were never observed after steady asphyxia. The pups displayed some abnormal behavior including shaking, twitches and Straub tail during the first 10 minutes of recovery. Thereafter, their behavior became indistinguishable from controls. In sharp contrast to this, intense behavioral seizures were seen in 7/15 pups shortly after intermittent asphyxia, with a median latency of 224 s [range: 178-310 s] from the end of the exposure (Fig. 2A and SVideo 1-2). Seizures of increasing severity (RSIII-RSV) occurred in succession. They commenced with forelimb clonus (RSIII) which was often coupled with rhythmic head-nodding. This was followed by clonus with loss of righting (RSIV), and the seizures ended after an episode of tonus-clonus with loss of righting in all 7/15 seizing animals (RSV). The median total duration of seizures was 110 s (range: 66-180 s), and all observed seizures terminated within about 8 min after the end of intermittent asphyxia. This was followed by a period (typically 40-180 s) of total immobility apart from respiratory movements, after which normal behavior gradually resumed. To our knowledge, this is the first description of a rodent model of human full-term birth asphyxia, in which robust seizures emerge after the termination of the insult, as observed in the Neonatal Intensive Care Unit (e.g. Lynch et al., 2012 and discussion).

**Fig 2.**
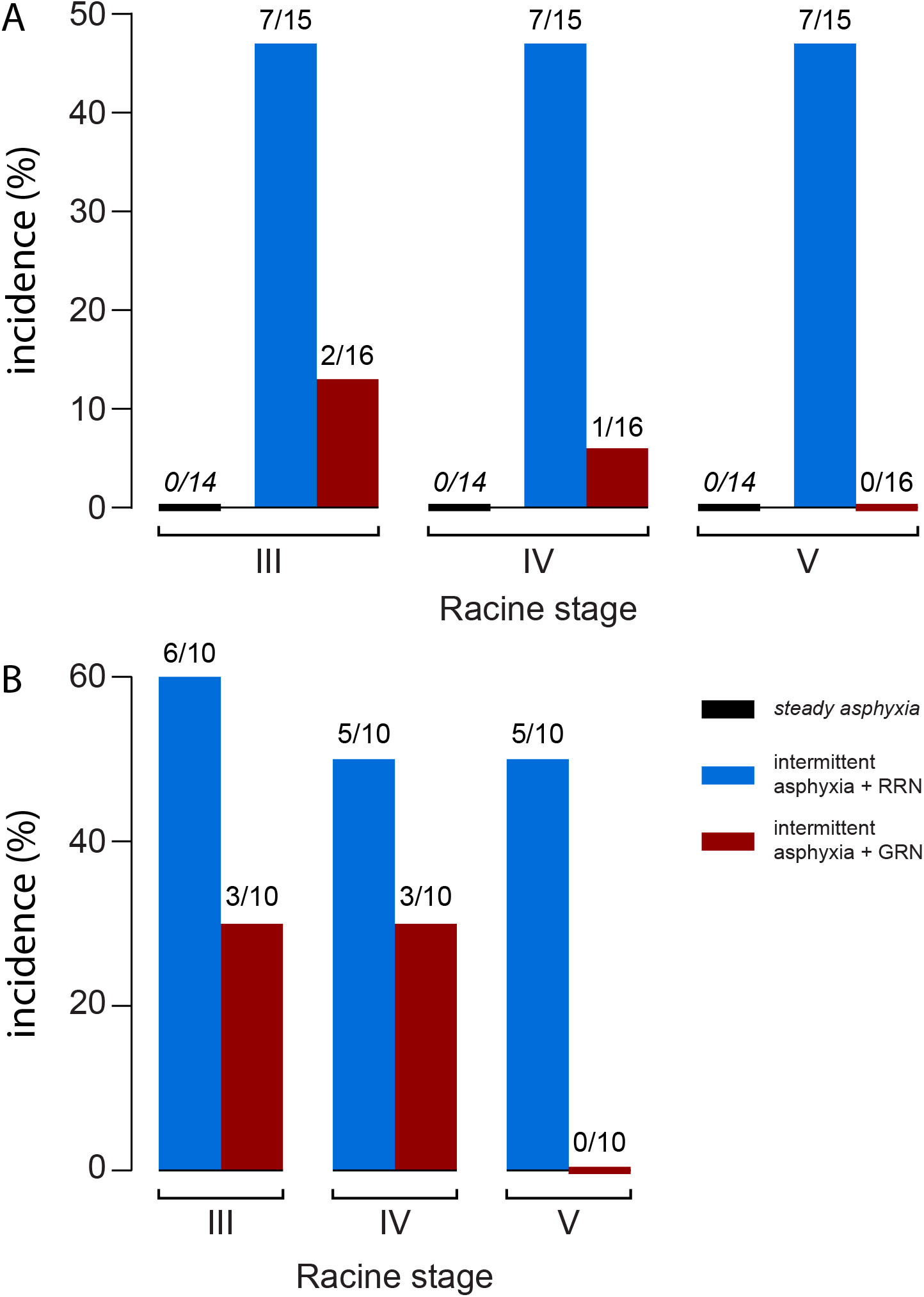
Incidence of Racine stage III-V behavioral seizures after experimental asphyxia. **(A)** Freely moving, non-instrumented rats were exposed to steady or intermittent asphyxia followed by rapid (RRN) or graded restoration of normocapnia (GRN). No seizures were seen after steady asphyxia, whereas they were triggered in 7/15 animals after intermittent asphyxia with RRN. GRN significantly decreased the proportion of animals with RSIII-IV seizures and abolished RSV seizures. **(B)** Similar to (A) but the animals had been implanted with epidural, cortical electrodes 24 h earlier (see Fig. 3 for electrocorticography). GRN abolished RSV seizures also in these experiments.

The severe acidosis of the brain that takes place during asphyxia shows a prompt recovery to the alkaline direction after the end of the exposure (see Bender et al., 2003; Pospelov et al., 2020; Remzso et al., 2020). There is evidence indicating that both the amplitude and the rate of alkaline recovery of brain tissue pH, boosts neuronal excitability and the triggering of seizures (Woodbury et al., 1957; Yoshioka et al., 1996). Thus, we examined the efficacy of Graded Restoration of Normocapnia (GRN) (Helmy et al., 2011), in suppressing seizures by exposing the rat pups to 5 % CO_2_ in air for 30 min immediately after intermittent asphyxia. Here, Rapid Restoration of Normocapnia (RRN) refers to the bulk of experiments in which the animals were promptly re-exposed to room air after asphyxia. Indeed, we found that GRN significantly reduced the occurrence of convulsive seizures compared to RRN (GRN 2/16 vs. RRN 7/15, p=0.0379 [Barnard’s test]; Fig. 2 and SVideo 3). Notably, none of the animals in the GRN group had RSV seizures (p=0.0016 compared to RRN). Thus, GRN reduced both seizure incidence and severity.

### Electrocorticographic and behavioral post-asphyxia seizures in freely moving rats

To examine the effects of intermittent asphyxia on neocortical activity patterns and the relationships between electrographic and behavioral seizures, electrocorticography (ECoG) was recorded at a frontal and a parietal site. Under control conditions, the ECoG was continuous and, characteristic of this age point, discrete bursts of activity were rarely observed (Cirelli and Tononi, 2015; Gramsbergen, 1976) (Fig. 3A, excerpt *a*). Immediately following the onset of intermittent asphyxia, the ECoG activity was strongly suppressed. Most of this suppression is caused by the hypercapnic acidosis as will be demonstrated below (Figs. 4 and 5). Nevertheless, during the initial phase of the asphyxia with 9 % of O_2_, about 25 % of the ECoG power persisted, while the subsequent fall to 5 % O_2_ led to a further decrease with hardly any detectable activity. These effects were constant during the three transitions from 9 % to 5 % O_2_ (Fig. 3A, lower panel and excerpt *b-c*).

**Fig 3.**
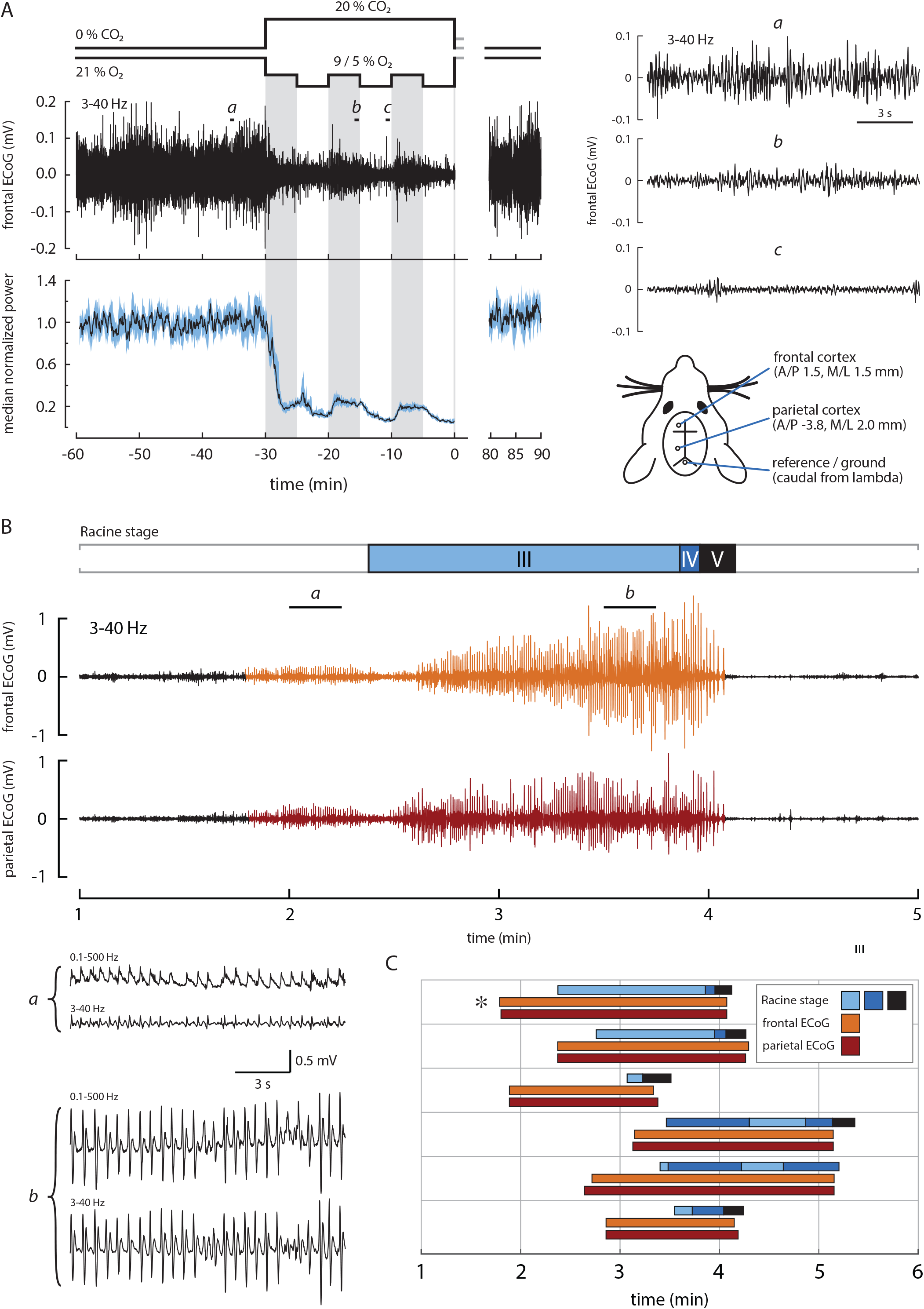
Electrographic and behavioral seizures emerge after but not during intermittent asphyxia. **(A)** Experimental design and the effect of asphyxia on cortical activity. The panels on the left show the gas application protocol (top), an example of a frontal electrocorticography (ECoG) recording (middle) and median power [95 % CI] of 18 frontal ECoG recordings (bottom). Excerpts a-c marked on the example ECoG are shown on the right. These data demonstrate a profound suppression of cortical activity during asphyxia. The placements of electrodes are shown in the illustration on the lower right corner. **(B)** An example of a post-asphyxia recording (asphyxia ends at time 0) showing the temporal relationship between behavioral and electrographic seizures. Top panel shows the behavioral seizure stages and middle and bottom panels show frontal and parietal ECoG, respectively. Excerpts a and b from the frontal ECoG displaying the early and the late parts of the seizures are shown in the lower left corner. We included two frequency bands to show the seizure activity appears without possible filtering artefacts. (C) Timeline representation of all ECoG (frontal and parietal) seizures and behavioral seizures in the RRN group. The recording marked by an asterisk is the same as in (B).

**Fig 4.**
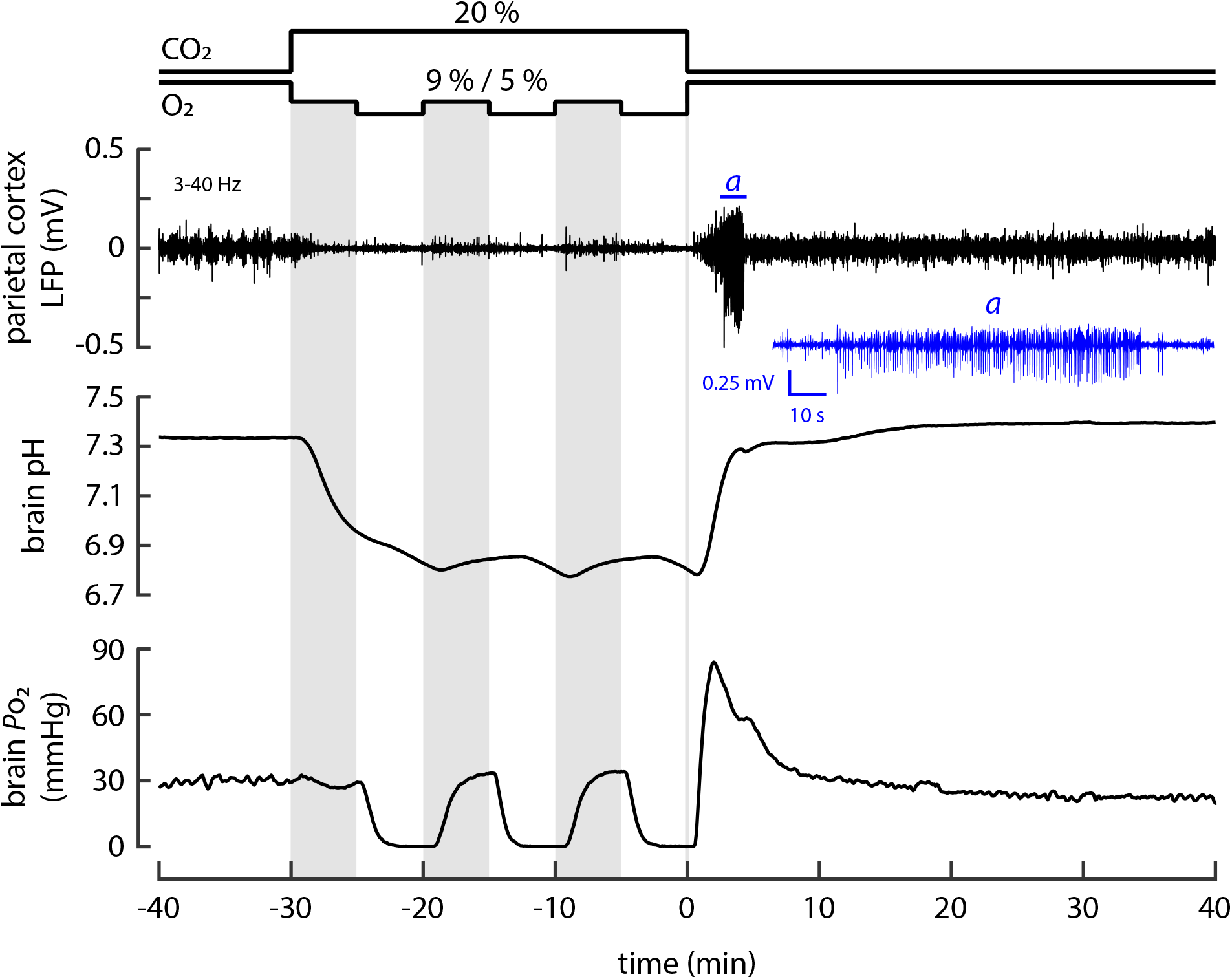
A sample recording with simultaneous monitoring of parietal cortex local field potentials (LFP), as well as brain pH and PO_2_. After the intermittent asphyxia, prominent seizure activity was seen in the LFP recording (top panel). Excerpt of the LFP trace (marked with a) including the seizure epoch is shown below in blue. Middle and bottom panels show the pH and PO_2_ recordings, respectively. To facilitate visual inspection, the three gray vertical columns mark the 9 % O_2_ bouts during the intermittent asphyxia protocol.

**Fig 5.**
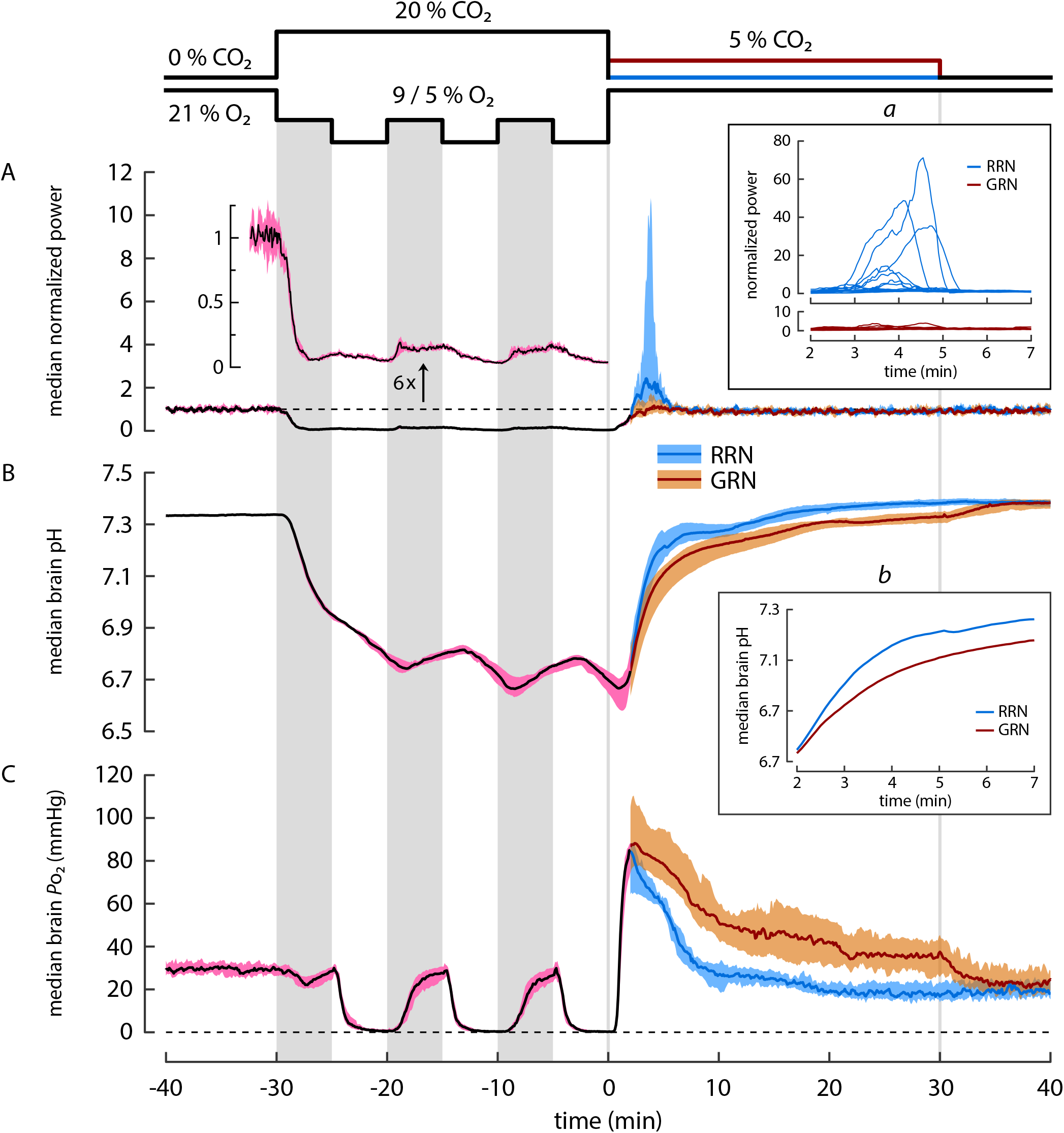
Changes in cortical activity (A) and in brain pH (B) and PO_2_ (C) in rat pups exposed to intermittent asphyxia, followed by rapid (RRN) or graded restoration of normocapnia (GRN). All values are median [95 % CI] except for panel A inset a. The three vertical grey columns on data panels mark the 9% O_2_ bouts during the intermittent protocol whereas the two vertical lines indicate the end of asphyxia and the end of GRN. Median data for the RRN and GRN protocols are plotted separately starting at 2 min after the end of asphyxia as the conditions before this time point are identical in the two protocols. **(A)** Power of local field potentials (LFPs) in the parietal cortex. The trace with a separate y-axis is a six-fold magnification of the LFP power and shows the profound suppression of cortical activity during asphyxia (cf. Fig. 4). After asphyxia, the activity recovers promptly. Following RRN, but not GRN, a period of hyperexcitability and seizures was seen, and it coincided with the rapid alkaline recovery of brain pH during a time window of 2 to 7 min after the end of asphyxia. The inset a shows individual LFP power traces during this period. **(B)** Brain pH decreases by 0.6 pH-units during the asphyxia and displays only minor modulation (0.12 pH-units) upon the changes between the 9% and 5% O_2_ levels. The alkaline recovery of brain pH after RRN is biphasic, and it is slowed down by GRN. The brain pH during the first recovery phase (2-7 min) is shown magnified in inset b. **(C)** Brain PO_2_ falls to apparent zero during the three periods of exposure to 5 % O_2_, but recovers to achieve a normoxic level when the ambient O_2_ level is increased to 9 % O_2_. After the asphyxia, an overshoot of PO_2_ is observed. GRN increases the duration but not the amplitude or rate of the PO_2_ overshoot.

In line with the purely behavioral observations in Fig. 2, we never observed electrographic seizure activity during the asphyxia exposure. In contrast, the ECoG recordings showed intense post-asphyxia seizure activity in 6 out of 10 freely-behaving animals which is in agreement with the seizure incidence in the non-instrumented animals. The electrographic seizures in the parietal cortex had a median latency of 150 [95 % CI: 108-188] s after the end of asphyxia. In all animals, the electrographic seizures appeared practically simultaneously at the recording sites in the parietal and the frontal cortex. The duration of the seizures in the frontal and the parietal cortex was comparable (pooled median duration 118 [95 % CI: 87-137] s) and most of the seizures consisted of spike trains with a 1.2-2.3 Hz discharge frequency (see Fig. 3B for an example) with increasing amplitudes towards the end of the seizure epoch. The ECoG bursts in the initial seizure period (Fig 3B, excerpt *a*) were monopolar (without the 3 Hz high-pass filtering), and we did not observe slow components (Fig 3B, excerpt *a-b*). The seizure termination was abrupt (delay from high amplitude spiking to complete cessation was 0-7 s), and it was followed by a strong suppression of the background ECoG. Thereafter, baseline activity gradually recovered.

The ECoG recordings demonstrated a consistent temporal overlap between the convulsive behavioral seizures and the electrographic seizures (Fig. 3B-C). The ECoG seizures preceded the convulsive seizures by 19 to 71 s. During this time, transient bouts of abnormal behavior, such as shaking, twitches and Straub tail, were observed. These aberrant behaviors were reminiscent of the non-convulsive RSI-II seizures, which were excluded from the present analyses (see Methods). The onset of the behavioral seizures and transition from one seizure stage to another was not marked by any obvious change in the ECoG (SVideo 4). In 4/6 cases, RSV seizures, which are considered to originate from the brainstem (Kellaway and Hrachovy, 1983; Pospelov et al., 2016), continued for an additional 3-13 s after the cortical seizures had terminated (Fig. 3C).

GRN decreased the post-asphyxia seizure incidence from 6/10 to 3/10 as seen in both behavior and ECoG. In agreement with the data on freely moving non-instrumented animals (Fig. 2A), RSV seizures were absent in the instrumented GRN group as well (Fig. 2B). Otherwise, the seizures were qualitatively similar to those observed after RRN. No behavioral or electrographic seizures were observed in the control ECoG group (n=7).

### Simultaneous recording of neocortical LFPs, brain pH and brain oxygen levels during intermittent asphyxia

While the data presented in Fig. 3 show that changes in ambient CO_2_ and O_2_ exert robust effects on neuronal excitability (from profound suppression to hyperexcitability during and after intermittent asphyxia, respectively), they provide no information on the contributions of the associated changes in brain pH and brain *P*o_2_. Therefore, we made simultaneous recordings of local-field potential (LFP) activity and pH as well as *P*o_2_ from the parietal cortex in head-fixed urethane-anesthetized rats.

The sample recording in Fig. 4 shows the quality of the raw data. Exposure to intermittent asphyxia (with the initial 9 % O_2_) caused a fast decline in brain pH, and the ongoing LFP activity, and the three shifts in ambient O_2_ from 9 % to 5 % produced a further, reversible suppression (cf. Figs. 3 and 5). A major decrease in *P*o_2_ was caused only by the three exposures to 5 % O_2_. A prompt increase in neuronal excitability took place leading to seizure activity within 2.8 min post-asphyxia in a manner comparable to what was seen in freely moving animals (Fig. 3). Moreover, the seizure pattern was similar, consisting of epileptiform spikes with a crescendo pattern.

### Suppression of LFP activity during intermittent asphyxia is mainly attributable to the fall in brain pH

Simultaneous recordings of LFP, pH and *P*o_2_ were made in a total of 29 animals exposed to intermittent asphyxia and followed by RRN (n=17) or GRN (n=12). During the first 5 min of intermittent asphyxia (with 9 % O_2_), the median value of brain pH (RRN and GRN groups combined) decreased rapidly from 7.33 to 6.95, with a further decline to 6.78 during the next 5 min in 5 % O_2_ (Fig. 5B). pH recovered only slightly (0.12 units; less than 20 % of the maximum acidosis) in response to the two subsequent 9 % O_2_ bouts, and dropped below 6.7 during the second and third 5 % O_2_ exposures, reaching a minimum of 6.67.

Median brain *P*o_2_ during the pre-asphyxia baseline was 30.0 mmHg. During the first 5 min period of intermittent asphyxia with 9 % O_2_, brain *P*o_2_ showed initially a small transient fall (nadir at 22 mmHg), but promptly recovered back to the baseline level, *indicating a fast re-establishment of brain normoxia* despite the large fall in ambient O_2_ (Fig. 5C). Notably, however, under these conditions, median LFP power was still suppressed to 10 % [95 % CI: 8.3 % – 11 %] from its baseline level, which is readily explained by the pronounced acidosis. A similar recovery to brain normoxia, associated with a strong suppression of LFP when compared to baseline (to 16 % [13 % – 19 %]), was observed during the two subsequent exposures to 9 % O_2_. In contrast to the brain normoxia achieved during the 9 % O_2_ exposures, all three epochs with 5 % O_2_ led to a decrease in brain *P*o_2_ to a level where virtually no free oxygen could be detected. Under these conditions, LFP activity was – consistent with the associated acid shift – further suppressed, to 4.1 % [3.3 % – 5.1 %] of the baseline power (see magnified power trace in Fig. 5A). Thus, all the above data indicate that the LFP power was suppressed by up to 90 % despite brain normoxia at 9 % ambient O_2_ because of the simultaneous hypercapnic acidosis.

### Dependence of the post-asphyxia increase in neuronal excitability and seizures on brain pH changes

After RRN, brain pH recovered in a biphasic manner, apparently reflecting the different elimination rates of CO_2_ and lactacidosis (Pospelov et al., 2020). During the initial, fast phase starting 2 min and ending 7 min after the asphyxia, brain pH increased from 6.75 to 7.26 (Fig. 5B and inset *b* therein). This was followed by an almost linear and much slower secondary recovery phase (cf. Pospelov et al., 2020) lasting until 15 min post-asphyxia. We did not observe an alkaline overshoot of the kind described in P6 rats with steady asphyxia induced by 9 % O_2_ and 20 % CO_2_ (Helmy et al., 2012, 2011), indicating that fast brain pH recovery in itself is sufficient in triggering seizures following asphyxia (see also Panaitescu et al., 2018). GRN reduced the rate of brain pH recovery after intermittent asphyxia. The fast phase of pH recovery, which in the RRN group took 5 min (see above), was slowed down by a factor of 2.6, to 13 min, by GRN (Fig. 5B inset *b*). The rate of pH recovery during the subsequent slower phase with GRN was closely similar to what was observed after RRN, but it took place at a more acidotic level with its start set by the slower rate of the preceding fast phase. The full recovery of brain pH following GRN had a third, final phase which was caused by the change from 5% ambient CO_2_ to room air, and was completed within 10 min.

Following RRN, there was a prompt increase in neuronal activity. In 12/16 animals this was followed by the emergence of recurring spike bursts (single burst duration <10 s) which in three cases transformed into prominent seizures (duration 94-117 s). The power maximum of individual LFP recordings was observed at around 3.5 – 4.5 min after the asphyxia and showed large variation ranging from 1.3-71 x [baseline power] (median 2.4 x; Fig. 5A and inset *a* therein). The low incidence of clear-cut seizure activity in experiments of the kind illustrated in Figs. 4 and 5 is readily explained by the use of urethane anesthesia (Cain et al., 1992). Regardless, a period of hyperexcitability, coinciding with the fast phase of pH recovery (2.0-7.0 min after the asphyxia), was consistently associated with RRN (Fig. 5A inset *a*). In contrast to this, in the GRN experiments the median LFP power returned to the baseline level promptly after the asphyxia with no significant overshoot. Accordingly, the median LFP power 2-7 min after the end of asphyxia was significantly higher in the RRN than in the GRN group (RRN 1.5 [95% CI: 1.1-3.3] vs. GRN 1.1 [95% CI: 0.75-1.5]; p=0.0130 [Mann-Whitney U-test]).

The present data indicates that, while seizures are generated only at pH levels relatively close to baseline (>7.0), it is not only the absolute pH but also *the rate of change of pH within a defined pH window* which is tightly linked to the triggering of seizures. This conclusion is supported by the astonishingly accurate temporal overlap of the major (RRN) and the minor (GRN) increases in LFP power seen during the 2-7 min window after the asphyxia (Fig. 5A inset *a*).

### Absence of effect of brain O_2_ on seizure generation after intermittent asphyxia

When the animals in the RRN group were re-exposed to room air after asphyxia, median brain *P*o_2_ recovered rapidly (max rate 140 mmHg/min) with an overshoot to 85 mmHg (cf. 30.0 mm Hg during baseline) at 2.0 min after the end of the intermittent asphyxia (Fig. 5C). Brain *P*o_2_ fell to the baseline level 6.6 min later and showed a transient decline to 16.4 mmHg 26 min after asphyxia. GRN had no effect on the rate (max 146 mmHg/min) or magnitude (peak 88 mmHg) of the brain *P*o_2_ overshoot. However, GRN increased the duration of the overshoot, and median *P*o_2_ remained elevated for the whole 30 min period. Thereafter, the RRN and GRN *P*o_2_ curves merged. It is noteworthy that brain *P*o_2_ in both the GRN and the RRN group is well above baseline at the time when seizures emerge.

## DISCUSSION

We demonstrate here for the first time behavioral convulsions and neocortical electrographic seizures in a rat model, that physiologically mimics BA in human full-term neonates. Consistent with the BA seizures seen in babies the seizures are observed after, but not during, the asphyxia insult (Lynch et al., 2012). Our direct and simultaneous measurements of neuronal activity and brain tissue *P*o_2_ show beyond any doubt that the post-asphyxia seizures are not triggered by brain hypoxia. Our intermittent asphyxia model, in which robust seizure activity was observed, satisfies the standard clinical criteria of BA, including a pronounced fall in blood pH from 7.5 to 6.9, associated with a significant metabolic acidosis as indicated by a decrease of 20.6 mM in base excess, and an increase in lactate from 1.1 to 8.6 mM. Moreover, the plasma level of the tell-tale endocrine biomarker of BA, copeptin (Evers and Wellmann, 2016; Kelen et al., 2017; Summanen et al., 2018), increased from 0.2 to 6 nM. As a whole, our data demonstrate that the present BA-seizure model is physiologically valid in that it preserves the organism’s innate responses to asphyxia and to the recovery thereof (see also Pospelov et al., 2020), which have been extensively studied in large-animal models of BA (Giussani, 2016; Lear et al., 2018) and obviously in human neonates as well (Evers and Wellmann, 2016; Lear et al., 2016; Vutskits, 2014).This point is of utmost importance because, as discussed below, the changes in the excitability of the brain, including the generation of seizures, are controlled by mechanisms that act not only within the brain, but also at the whole-organism level.

### Hypercapnia as an endogenous seizure-suppressing factor during the asphyxia-linked hypoxia

Asphyxia is, by definition, a combination of systemic hypercapnia and hypoxia. The associated hypercapnic acidosis of the brain leads to suppression of neuronal activity as is presently evident in the ECoG and LFP recordings in Figs 3-5 (Lee et al., 1996; Lennox, 1936; Mitchell and Grubbs, 1956; Pollock, 1949; Tolner et al., 2011; Ziemann et al., 2008), while alkalosis has the opposite effect (Lee et al., 1996; Lennox, 1936; Nasreddine et al., 2020; Schuchmann et al., 2011, 2006). Indeed, a wide variety of neuronal ion channels, which control intrinsic neuronal excitability and synaptic signaling, show a high and functionally synergistic sensitivity to pH (Pasternack et al., 1996; Spray et al., 1981; Tombaugh and Somjen, 1997; Traynelis and Cull-Candy, 1990; Wemmie et al., 2013; Wilkins et al., 2005). Thus, the suppression of excitability by acidosis during asphyxia is deeply rooted in mammalian evolution (see Ruusuvuori and Kaila, 2014). In line with this, the behavioral and neocortical seizures in our model are not triggered during, but only after the BA-mimicking insult.

In stark contrast to the above, in both non-invasive and invasive models of BA and/or HIE in which rat and mouse pups are exposed to pure hypoxia (4-8 % O_2_ in N_2_) (Greggio et al., 2009; Jensen et al., 1991; Rakhade et al., 2011; Rodriguez-Alvarez et al., 2015; Sampath et al., 2014; Zanelli et al., 2014; Zayachkivsky et al., 2015), seizures are triggered already *during* the hypoxia insult. Strikingly, the highly artificial condition of pure hypoxia leads to a brain *alkalosis* (Mitsufuji et al., 1995; Pospelov et al., 2020), which is known to produce an increase in the excitability as well as seizures especially in the immature brain (Miley and Forster, 1977; Ruusuvuori et al., 2010; Schuchmann et al., 2011, 2006; Wirrell et al., 1996). Here, it is worth emphasizing that peri- and neonatal rodents or other mammals are, under non-experimental conditions, never exposed to anything that could be called “birth hypoxia”.

### Post-asphyxia seizures are not strictly related to the preceding hypoxic load

Our data on blood acid-base parameters (Fig. 1) indicate that the hypoxic load is near-identical in the intermittent vs. the steady asphyxia protocol used presently. Thus, the post-asphyxia seizure generation is not strictly related to the magnitude of the hypoxic load. While the intermittent asphyxia protocol resulted in severe behavioral post-asphyxia seizures (RSIII-V) in about half of the animals tested, we never observed seizures following steady asphyxia. A possible scenario that might account for the efficacy of intermittent asphyxia in triggering seizures is that mechanisms, which have been shown to promote anoxic/hypoxic LTP *in vitro* (Di Filippo et al., 2008; Hsu and Huang, 1997; Quintana et al., 2015), are likely to be activated during the 5 % O_2_ bouts of intermittent asphyxia. The enhanced network activity during the periods with 9 % O_2_ (and transient brain normoxia) would then lead to further activity-dependent potentiation of excitatory connections. Finally, during the recovery from asphyxia, the suppressing effect of the hypercapnic acidosis is quickly removed, unmasking the enhanced excitability, which results in seizure generation. Here, it is worth pointing out that while much work on rodent models has focused on neuronal molecules and signaling mechanism that are affected as a *consequence* of seizures (Lippman-Bell et al., 2016; Rakhade et al., 2011; Sun et al., 2013; Wang et al., 2011; Zhou et al., 2015), next to nothing is known about the *proximate causes in vivo* which render the neonate’s brain prone to seizures after BA.

### Post-asphyxia seizure generation shows a steep dependence on the rate of recovery of brain pH but not oxygen

The convulsive behavioral seizures after intermittent asphyxia were closely paralleled by neocortical seizures as demonstrated by ECoG recordings. Careful reviewing of the video-ECoG recordings and the movement-transducer signal showed only minor contamination of the ECoG by movement artefact even during intense convulsions (see e.g. SVideo 4, section 2:20-2:26). The seizures were detected in the frontal and the parietal cortex practically simultaneously indicating that the post-asphyxia seizures spread rapidly over large cortical areas (Fig. 3). This observation also implies that our 2-site recording is reliable for neocortical seizure detection in the present model. The electrographic seizures had initially a low amplitude (Figs. 3-4) which increased towards the end of the discharge. The crescendo and the discharge frequency resembled those observed in rat pups during hyperthermia-induced brain alkalosis (Ruusuvuori et al., 2013; Schuchmann et al., 2006). In line with this, both the incidence and severity of the behavioral as well as the electrographic seizures seen after RRN were strongly reduced by GRN (Figs. 3 and 5), which in the present work was achieved by application of 5 % CO_2_ in room air immediately after the asphyxia.

The steep dependence of post-asphyxia hyperexcitability and seizures on recovery of brain pH (but not O_2_) was directly demonstrated in simultaneous recordings of cortical pH, *P*o_2_ and LFP activity done under light urethane anesthesia. In these experiments, a prominent period of post-asphyxia hyperexcitability took place, during which some animals (3/16) developed frank seizures (Fig. 5). The time window of hyperexcitability and seizures coincided with the fast post-asphyxia recovery of brain pH and, again, GRN led to a near-complete abolishment of post-asphyxia hyperexcitability. Notably, the rate of recovery and overshoot of brain *P*o_2_ were similar following GRN and RRN, demonstrating a lack of contribution of the *P*o_2_ changes to seizure suppression by GRN. It is obvious from the data in Figs 3 and 5 that a much briefer duration (around 10 min, given the seizure time window of 2-7 min post-asphyxia) than the pre-set time of 30 min for the present experiments would have been sufficient for the anticonvulsant action of GRN.

The pH-related seizure episode (about 2 min) in our model is obviously much shorter than in human neonates. But here, brain and body size do matter: while the recovery of systemic and brain pH takes only 15 min after the asphyxia in the rat pups, in asphyxiated human neonates the recovery of systemic pH can last up to 20 h. This time window overlaps with seizure onset in the human neonate (median 17.1 h [inter-quartile range 12.5–21.3] in Lynch et al., 2012). While no data are available for post-asphyxia pH recovery in the human brain, a point of reference is provided by the piglet in which post-asphyxia blood pH and brain pH recovery takes 40-120 min (Bender et al., 2003; Remzso et al., 2020). For comparison the body weights of a P11 rat, a newborn piglet and a newborn human are about 21, 1200 and 2900 g and the brain weights 1, 33 and 370 g, respectively (Bandeira et al., 2009; Dekaban and Sadowsky, 1978; Minervini et al., 2016), Thus, the duration of pH recovery and BA seizures in the present rat model are, in meaningful allometric manner, consistent with those in larger mammals.

### Conclusions

The present work shows that post-asphyxia changes in brain pH have a pronounced effect on seizure susceptibility. We do not, of course, argue that brain pH changes would be the most important factor in all kinds of post-asphyxia seizures in human neonates. A number of mechanisms triggered by e.g. energy-metabolic compromise and neuroinflammation are likely to contribute (Schiering et al., 2014). However, the dependence of neuronal excitability on brain pH is particularly steep in the immature brain (Ruusuvuori et al., 2010). Thus, it is reasonable to assume that the large pH changes, which are bound to take place during and following birth asphyxia in human neonates (see Uria-Avellanal and Robertson, 2014), are likely to have a major influence on the generation and properties of post-asphyxia seizures.

Based on a number of criteria described above, our work calls into question the translational validity of rodent BA models, which are based on exposure of neonate animals to pure hypoxia. Indeed, by far most of the translational impact of preclinical work has come from large-animal models of BA/HIE, where the experimental protocols typically induce a state of asphyxia, not hypoxia (Giussani, 2016; Hassell et al., 2015; Koehler et al., 2018; Lear et al., 2018). Moreover, large-animal models have numerous advantages with regard to instrumentation in studies on adaptive and pathophysiological mechanisms associated with BA (Drury et al., 2015; Hassell et al., 2015; Saugstad et al., 2019; Shim et al., 2020). On the other hand, standard laboratory rodents offer the important potential for implementing the vast array of current neurobiological techniques which are available for both basic and translational work on the pathophysiology, neurobiology as well as behavioral outcome of BA and HIE. We are confident that physiologically-valid rodent models of the kind described presently will make a significant contribution to the research in this highly important field.

## Supporting information

SVideo1

SVideo2

SVideo3

SVideo4

## ACKNOWLEDGEMENTS

We thank Merle Kampura for assistance in the molecular biological analyses, and Maria Partanen, Madara Snepere and Ann-Christine Aho for the breeding and maintaining of the experimental animals.

