## Supplement captions for "A physiologically-validated rat model of term birth asphyxia with seizure generation after, not during, brain hypoxia"

**SVideo 1:** Behavioral video of a P11 rat pup showing intense seizures during recovery from intermittent asphyxia. Video begins 3 min after the end of asphyxia and rapid restoration of normocapnia (RRN). Seizure stages are shown on the left. All rat pups in the supplementary videos are P11-12.

**SVideo 2:** Behavioral video of the recovery of a second animal from the intermittent asphyxia + RRN group (cf. SVideo 1). Video begins 3 min and ends 5.5 min after asphyxia. The present work as a whole suggest that seizure probability is highest within this timeframe. However, no seizures were observed during, before or after this time period in this example.

**SVideo 3:** Behavior during graded restoration of normocapnia, 3 – 5.5 min after the end of intermittent asphyxia. No seizures were observed during, before or after this time period.

**SVideo 4:** Video-electrocorticography (ECoG) showing intense seizures in a rat pup that was exposed to intermittent asphyxia followed by RRN. Video begins 90 s after asphyxia. From bottom up the channels are piezo, parietal ECoG and frontal ECoG. The top three state channels mark seizures in the parietal ECoG, the frontal ECoG and behavioural seizures. Racine stages III-V are marked with blue, green and teal, respectively, in the topmost channel.
